# Probabilistic method corrects previously uncharacterized Hi-C artifact

**DOI:** 10.1101/2020.10.07.325332

**Authors:** Yihang Shen, Carl Kingsford

## Abstract

Three-dimensional chromosomal structure plays an important role in gene regulation. Chromosome conformation capture techniques, especially the high-throughput, sequencing-based technique Hi-C, provide new insights on spatial architectures of chromosomes. However, Hi-C data contains artifacts and systemic biases that substantially influence subsequent analysis. Computational models have been developed to address these biases explicitly, however, it is difficult to enumerate and eliminate all the biases in models. Other models are designed to correct biases implicitly, but they will also be invalid in some situations such as copy number variations. We characterize a new kind of artifact in Hi-C data. We find that this artifact is caused by incorrect alignment of Hi-C reads against approximate repeat regions and can lead to erroneous chromatin contact signals. The artifact cannot be corrected by current Hi-C correction methods. We design a probabilistic method and develop a new Hi-C processing pipeline by integrating our probabilistic method with the HiC-Pro pipeline. We find that the new pipeline can remove this new artifact effectively, while preserving important features of the original Hi-C matrices.

## 1 Introduction

The spatial arrangement of chromosomes is related to gene regulation processes such as transcription [Fraser and Bickmore, 2007] and epigenetics modifications [Grewal and Moazed, 2003]. Nuclear organization and its role in gene regulation have been studied broadly in recent years [Van Bortle and Corces, 2012, Rinn and Guttman, 2014]. Chromosome conformation capture (3C) [Dekker et al., 2002] and its derivatives [Simonis et al., 2006, Lieberman-Aiden et al., 2009] is a class of methods that can detect spatial contacts of chromosomes, providing new insights on the three-dimensional structures for chromosomes. Hi-C [Lieberman-Aiden et al., 2009], a widely used high throughput 3C-based method, outputs millions or even billions of read pairs, each representing one contact between two genomic loci. Each end of a read pair aligns to the reference genome and then is assigned into a genomic bin, an interval with fixed length such as 10 kilobase. This produces a two-dimensional contact matrix, called a Hi-C matrix. Each entry of the matrix represents the amount of contacts between two genomic bins *A* and *B*, where the value of an entry is the number of read pairs that have one end in *A* and the other in *B*.

Raw Hi-C read counts are affected by various kinds of biases. Imakaev et al. [2012] divided these biases into two categories, technical biases such as sequencing mistakes and mapping errors, and biological factors such as inherent properties of chromosomes. Yaffe and Tanay [2011] pointed out several systemic biases, such as the length of restriction fragments, GC content, and mappability (genomic uniqueness) of the fragment ends. Because these biases will affect the accuracy of the downstream analysis of chromatin structures, several Hi-C correction methods have been developed. Lieberman-Aiden et al. [2009] used a method called Vanilla coverage normalization (VC) to correct for bias, where each entry in a Hi-C contact matrix is divided by a row-specific normalization term and subsequently divided by a column-specific normalization term, generating a normalized contact matrix. Yaffe and Tanay [2011] developed a probabilistic method to model and correct systemic biases explicitly. Hu et al. [2012] further proposed a simplified, Poisson-based probabilistic model to enhance the efficiency as well as reproducibility of the normalization step. Imakaev et al. [2012] factorized values of entries in Hi-C matrices into the multiplication of contact probabilities and bias factors and then developed a method called iterative correction and eigenvector decomposition (ICE) to remove biases implicitly. Rao et al. [2014] used the KR matrix-balancing algorithm [Knight and Ruiz, 2013] to normalize Hi-C matrices and remove biases. Other correction work focused on normalizing special chromatin properties such as copy number variation in cancer cells [Vidal et al., 2018, Servant et al., 2018, Khalil et al., 2019].

In this work, we characterize a new kind of bias of Hi-C data. Our work shows that this bias is generated from the read alignment step, and can lead to significant, long-range chromatin contact signals. We present evidence showing that many signals of long-range contacts in Hi-C matrices are in fact artifacts. We find that these incorrect contact signals are conserved in Hi-C data from different cell types in our test regardless of Hi-C protocols or data processing pipelines used. Because of this highly conserved property, these contacts may be identified by mistake as important chromatin structures such as loops and may be mistakenly considered to play essential roles in some gene regulation. We find that the new bias cannot be corrected by current Hi-C correction methods. Therefore, we develop a new correction method by using probabilistic models. In our method, we use the empirical genomic distance distribution of read pairs as prior probability and calculate posterior probabilities of alignment locations of read pairs by integrating the information of base calling errors and genetic variations. Results show that our new method can filter these erroneous contacts, decreasing the false-positive ratio of chromatin loop calling, without destroying other typical features of Hi-C matrices such as TAD boundaries.

## 2 Identification of erroneous chromatin contacts

### 2.1 Data

We analyzed four non-cancer cell lines, lung fibroblast (IMR90), mammary epithelial (HMEC), umbilical vein endothelial (HUVEC) and epidermal keratinocyte (NHEK) from Rao et al. [2014]. All of the Hi-C data were downloaded as raw reads, and then processed by HiC-Pro pipeline [Servant et al., 2015], in which Bowtie2 [Langmead and Salzberg, 2012] was used for read alignment. Hi-C contact matrices were normalized by several normalization methods: ICE [Imakaev et al., 2012], KR [Rao et al., 2014] and VC [Lieberman-Aiden et al., 2009]. We downloaded CTCF ChIP-Seq peaks of all four cell types from the ENCODE project website [ENCODE Project Consortium et al., 2012]. We also downloaded RAD21 ChIP-Seq peaks and SMC3 ChIP-Seq peaks of IMR90 from ENCODE. We downloaded super-enhancers of these four cell types from SEdb database [Jiang et al., 2019]. Juicer tools [Durand et al., 2016] and Integrative Genomics Viewer [Robinson et al., 2011] were also used for visualization and downstream analysis.

### 2.2 Highly conserved, long-range contacts in Hi-C

We used Fit-Hi-C [Ay et al., 2014], a package that can identify statistically significant intra-chromosomal contacts, to detect significant contacts from 10 kilobase resolution Hi-C data of all four cell types with additional requirement that contacts had genomic distance ≥ 50kb. Here, one significant contact means one entry in a Hi-C matrix that has a significantly high value, and 10 kilobase is the length of one genomic bin. We detected 218,028 significant contacts in IMR90, 18,410 in HUVEC, 20,917 in NHEK, and 30,631 in HMEC (read depth of IMR90 are significantly higher than other three cell types, so it is reasonable to detect much more contacts in IMR90). For each cell type, around 5% of these contacts are long-range, having genomic distance > 2Mb (2,148 in HMEC, 8,633 in IMR90, 1,269 in HUVEC, and 2,749 in NHEK). Further comparison showed that 4002 contacts are conserved among all four cell types, and 416 of them are long-range (genomic distance > 2Mb). In general, long-range chromatin contacts are rare in Hi-C matrices [Rao et al., 2014], therefore we investigated the mechanisms that cause these highly-conserved, statistically significant contacts. Since contacts in chromosomes are highly related to CTCF motifs [Rao et al., 2014], we investigated the relationship between Chip-seq CTCF peaks and these contacts. The 4002 conserved contacts were divided into two categories: contacts with genomic distance ≤ 2Mb or > 2Mb. We found that among all these four cell types, less than 10% of the contacts with genomic distance between 50kb and 2Mb have no CTCF peaks in either of the bins, while more than 65% contacts in the second category contain no CTCF signals (Figure S1). Considerable differences in terms of the existence of CTCF peaks suggest that CTCF may not the main reason of forming contacts in the latter category. Rao et al. [2017] reported that super-enhancers (SEs) played an important role in the formation of cohesin-independent loops, which is another mechanism of forming long-range contacts. Therefore, we checked the existence of SEs in the conserved contacts with genomic distance > 2Mb. We found that around 85% of them did not overlap with known super-enhancer locations in either of two bins (IMR90:354/416, NHEK:399/416, HUVEC:365/416, HMEC:313/416). These results suggest that conserved contacts with genomic distance > 2Mb may be more likely to be technical artifacts, not real contacts in chromosomes.

However, signals representing these long-range contacts clearly exist in heatmaps of Hi-C matrices. A representative example can be seen in Figure 1. In chromosome 2, Fit-Hi-C detected 16 significant and conserved contacts in the intersection region with two intervals 87300kb-88100kb and 111900kb-112700kb. The genomic distance between these two intervals is 24 Mb, around 10% of the length of chromosome 2, and most of these 16 contacts (14/16) contain no CTCF peaks on either of two bins in at least one cell type. However, these contacts exist in heatmaps of non-normalized Hi-C matrices among all four cell types (Figure 1), and their shapes are highly similar. We also tried four commonly used correction methods, KR, ICE, VC and VC_SQRT, but none of them remove these contacts (Figure S2). We further found that these contacts exist in Hi-C heatmaps generated from different Hi-C protocols (Figure S3), different read alignment software or post-processing pipelines (Figure S4, Figure S5), and different reference genome assembly (Figure S6). The results suggest that these long-range contacts are not generated by a specific protocol or pipeline, and previous correction methods are not able to correct for them.

**Figure 1:**
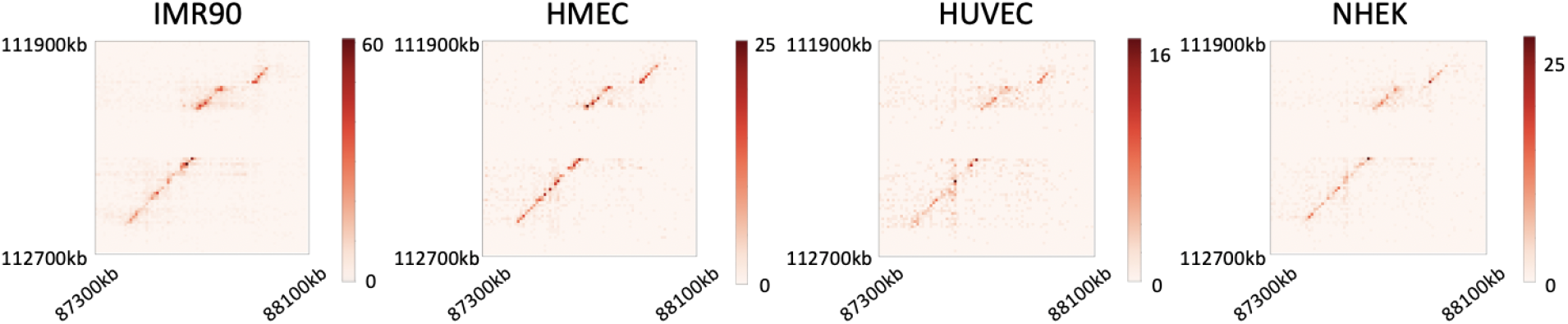
Representative example of a heatmap region containing highly conserved long-range contacts, i.e. two genomic intervals: 87300kb-88100kb and 111900kb-112700kb in chromosome 2. Resolution:10 kb. Heatmaps are from the non-normalized Hi-C matrices of all the four cell types.

### 2.3 Long-range contacts are artifacts from read alignment

Further experiments suggest that these long-range contacts are artifact and they are generated from incorrect read alignment. First, for each cell type we extracted all reads from long-range contacts (≥ 2Mb apart) which do not contain CTCF peaks. We then compared the distribution of mapping quality values (MAPQ) of these reads with the MAPQ distribution of all reads from one cell type, which we called reference reads. We found that the MAPQ value of reference reads is significantly higher than that from long-range contacts (*P* < 3 × 10^−308^, one-sided Mann–Whitney U test; Figure 2a), and a low MAPQ value suggests that one read could be incorrectly aligned.

**Figure 2:**
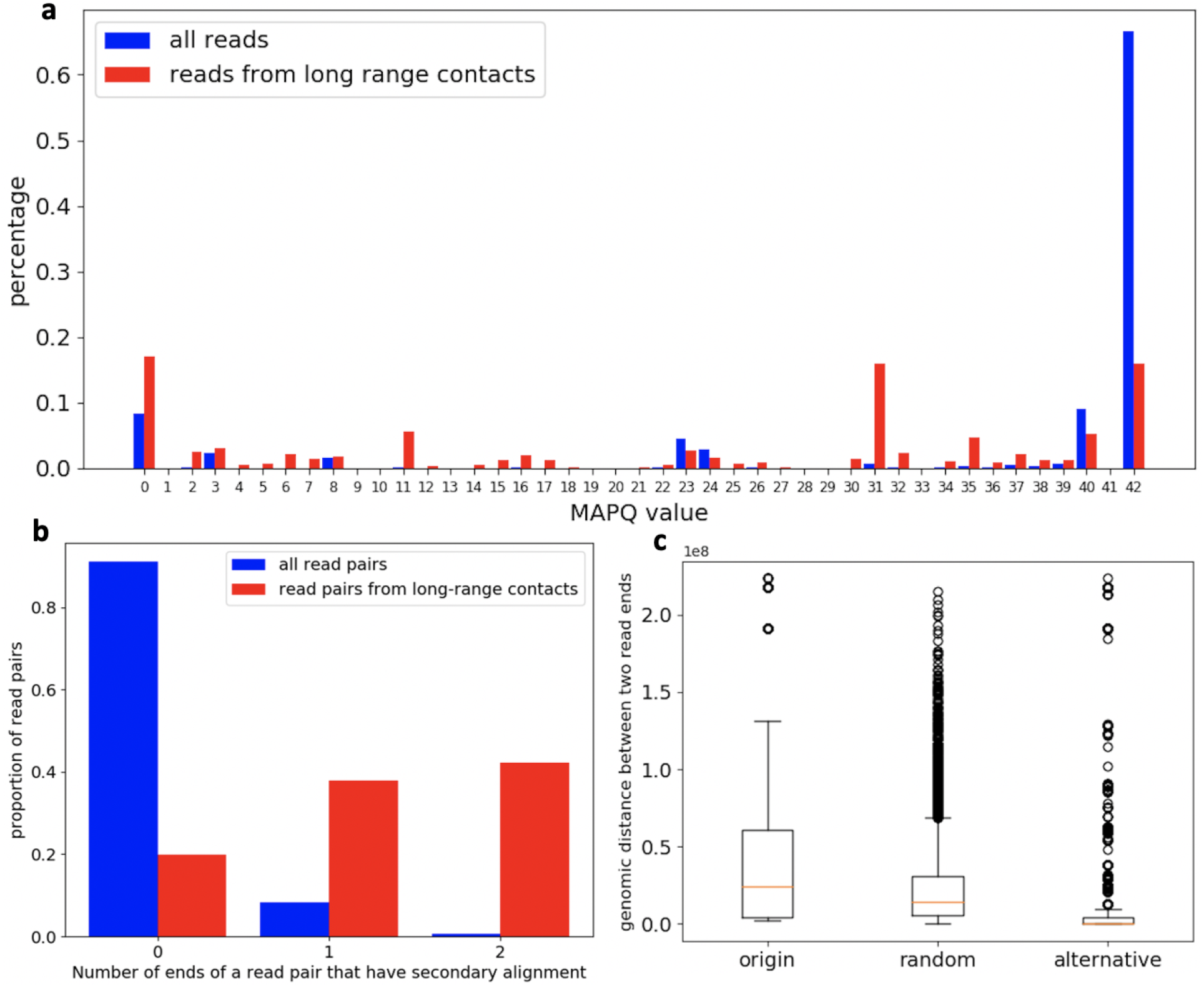
Evidence showing that long-range contacts are likely artifacts from read alignment. In this figure, data and results are from cell type NHEK. (a) The MAPQ distribution of all reads (reference reads) and reads from long-range contacts. (b) The proportion of all read pairs and read pairs from long-range contacts with three categories: neither of two ends in one read pair has secondary alignment position (the left column), one of them has secondary alignment positions (the middle column) and both of two ends have secondary alignment positions (the right column). (c) Box plots showing the distribution of genomic distance from original alignments of reads in long-range contacts (the column notated as “origin”), from random sampling (the column notated as “random”) and from possible minimal genomic distance choices according to secondary alignment results (the column notated as “alternative”).

If one read is aligned incorrectly, the correct position likely has a highly similar sequence to the incorrect one. Although the actual position does not get the highest mapping score at the read alignment step, it should be a near-optimal choice. Therefore, we investigated how many read pairs from long-range contacts that have secondary alignment choices reported by Bowtie2. The proportion of read pairs from long-range contacts that have secondary alignment positions is significantly higher than that from reference reads (P < 3 × 10^−308^, Chi-Square test) (Figure 2b), which suggests that most of these reads are from repeat regions.

We then investigated the locations of these secondary choices. We remapped the reads from long-range contacts that have secondary alignments by Bowtie2 again. Parameters were the same as before except that this time the top 2 alignments were reported for each read. After remapping, each read pair had 2 × 2 possible location pairs. We chose the location pair (*p*_1_, *p*_2_), such that *p*_1_ and *p*_2_ were from the same chromosome and meanwhile the genomic distance of (*p*_1_, *p*_2_) was the smallest among all the 4 possible choices. Hence the genomic distance of (*p*_1_, *p*_2_) can be regarded as the minimal possible intra-chromosomal genomic distance of one read pair (*r*_1_, *r*_2_). We also randomly sampled the minimal intra-chromosomal genomic distance as the background distribution. We found that most read pairs from long-range contacts have alternative location pairs with significantly smaller genomic distance than the origin of mapping and also than random sampling (Figure 2c). For example, for contacts in the intersection region with intervals 87740kb-87750kb and 112260kb-112270kb in chromosome 2, most reads that are originally mapped to 87740kb-87750kb have secondary alignments located in the interval 112257kb-112268kb, which is close to the other interval of the contact, 112260kb-112270kb. Therefore, it is likely that these read pairs are from two close genomic locations in reality but are aligned to two distant regions because of the sequence similarity between these two regions.

### 2.4 Repeats in chromosomes lead to erroneous long-range loops

Figure 3 shows the schematic diagram to illustrate a possible mechanism for the formation of these erroneous long-range contacts. Two repeats are shown in one chromosome. They have differences in some of the positions. For example, repeat 1 has base ‘A’ while repeat 2 has ‘T’ in the corresponding position. Given a Hi-C read pair, if one read end (or both two ends) is from repeat 1, and because of some variations of samples such as SNPs or incorrectness of the reference, it can be mapped to the other repeat, repeat 2. Long-range contacts then appear since repeat 2 is far away from the location that the other read belongs to.

**Figure 3:**
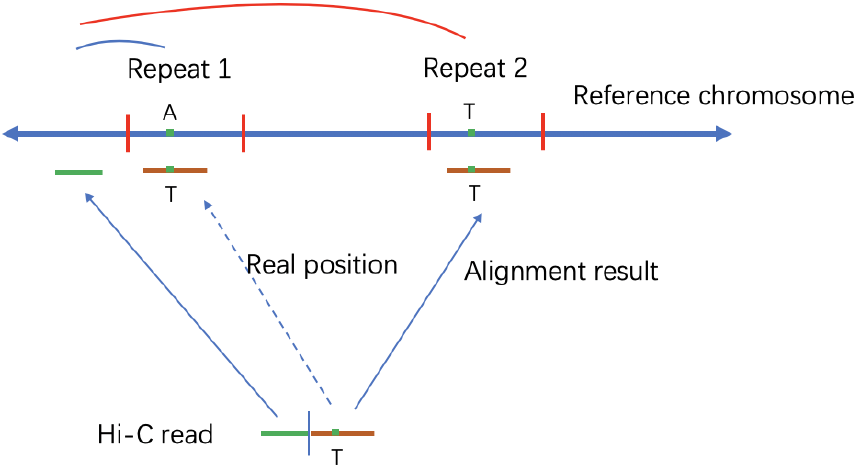
Illustration of the formation of erroneous long range contacts. Repeat 1 and 2 are two repeats from the same reference chromosome with a little differences, for example repeat 1 has base ‘A’ while repeat 2 has base ‘T’ in the same location. A Hi-C read pair with genomic variations such as SNPs may be mapped to the position of another repeat.

Rao et al. [2014] discussed briefly Hi-C contacts in repetitive regions, and they filtered multi-aligned reads out by removing all reads with MAPQ=0 (section VI.a.4.iii of the Extended Experimental Procedures). However, most reads from long-range contacts described above are not multi-mapped reads, instead they are the reads that have unique “best” alignments, although also have reasonable secondary alignment choices, and less than 20% of them have MAPQ value equals to 0 (Figure 2a). In fact, Figure S7 shows that simply filtering out reads with MAPQ=0 is not able to solve the problem of artifactual long-range contacts effectively.

## 3 Method

### 3.1 Probabilistic method for long-range contacts correction

We propose a probabilistic method to correct these artifactual long-range contacts. Given a Hi-C read pair (*r*_1_, *r*_2_), our goal is to find two sequences (*p*_1_, *p*_2_) from the reference genome (denoted as *X*) such that:

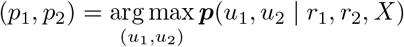

where (*u*_1_, *u*_2_) are two sequences from the reference. Here ***p***(*u*_1_, *u*_2_ | *r*_1_, *r*_2_, *X*) is the conditional probability of two sequences given one read pair and the reference. According to the Bayes’ rule, we have:

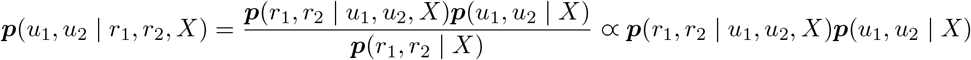

If we further assume that having reads from one genomic location is independent to having reads from the other location, we can factor ***p***(*r*_1_, *r*_2_ | *u*_1_, *u*_2_, *X*):

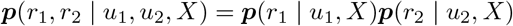

Then we have:

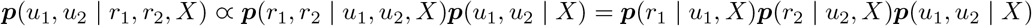

Furthermore, we assume that the prior ***p***(*u*_1_, *u*_2_ | *X*) is only related to the genomic distance between two sequences, here the sequence genomic distance is defined as the genomic distance between the first bases of these two sequences. We have ***p***(*u*_1_, *u*_2_ | *X*) = *f* (*D* | *X*) where *f* is the probability density function of genomic distance *D* = |*u*_1_ – *u*_2_|. (Note that our model only considers intra-chromosomal contacts since it is hard to build genomic distance distributions on inter-chromosomal contacts accurately.)

When estimating ***p***(*r* | *u, X*), we assume that all bases of one read are independent. Denote *r*[*i*] as the i-th base in the read *r*, denote *u*[*i*] as the i-th base in the sequence *u* from the reference, we can then factor **p**(*r* | *u, X*) as:

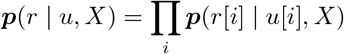

If we assume that there are no genetic variations between the samples and the reference genome, an assumption we will relax later, then the incorrect matching between the base from the read and the base from the reference is solely from the base calling error. Define the base calling error of *r*[*i*] as:

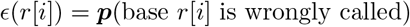

We have:

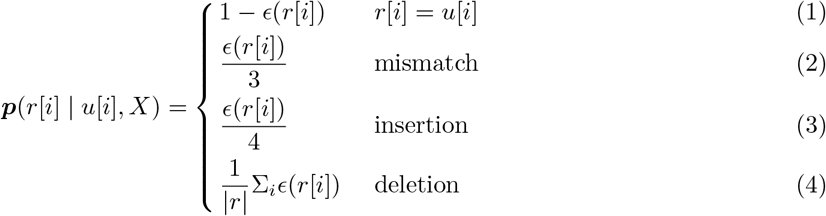

Equation (1) represents the probability of having a base that matches the reference, (2) is the probability of having a mismatch. Besides mismatch cases, here we also use empirical estimates of ***p***(*r*[*i*] | *u*[*i*], *X*) under insertion and deletion cases. For the insertion case (3), we simply divided *ϵ*(*r*[*i*]) by 4 since all the four kinds of bases are “wrong”, and for the deletion case (4), since the base of the read at this position is missing, we do not have any information about *ϵ*(*r*[*i*]), hence we use the average of *ϵ* among all the bases in the read *r* to estimate the deletion error, where |*r*| is the length of the read.

It is likely that the issue of long-range contacts comes from genetic variations instead of just base calling errors since many reads from the same genome region are mapped to incorrect positions. Therefore, we further integrate the information of genetic variations into our probabilistic model. Suppose SNPs are detected at position u[i], we denoted the probability of having a specific nucleotide at this position as: ***p***_*B*_ = ***p***(*B* | *u*[*i*], *X*) where *B* ∈{“A”,”T”,”C”,”G”}, which can be calculated by genotype calling. We defined the probability of having base *r*[*i*] from *u*[*i*] under genetic variants, ***p***_*GV*_(*r*[*i*] | *u*[*i*], *X*), as 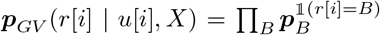 where 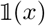 represents the indicator function. Therefore, the calculation of ***p***(*r* | *u, X*) can be divided into two parts, bases with or without genetic variants:

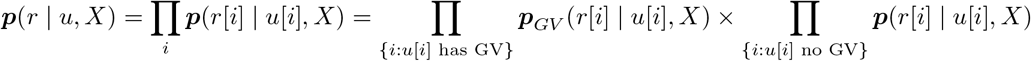

For each read pair (*r*_1_, *r*_2_), it is computationally infeasible to consider all the sequence pairs (*u*_1_, *u*_2_) from the reference and compute ***p***(*u*_1_, *u*_2_ | *r*_1_, *r*_2_,*X*). The read alignment step provides a set of most possible alignment choices (secondary alignments) for each read pair, denoted as *S*_1_ = {*u*_11_,…, *u*_1*i*_} for *r*_1_ and *S*_2_ = {*u*_21_,…, *u*_2*j*_ } for *r*_2_, and we only consider pairs from these two sets when calculating posterior probabilities. That is, we want to find two sequences such that:

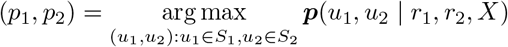

### 3.2 Combination with HiC-Pro pipeline

We integrate our repeat correction step with HiC-Pro, a widely used Hi-C processing pipeline, and call this LLC-HiC (long loop correction HiC). Figure 4a shows the workflow of LLC-HiC, which keeps all four main modules of HiC-Pro (blue part) and adds our correction method, Long Range Contact Correction module between the Read Alignment module and the Valid Pairs Detection module. Note that only intra-chromosomal interactions will be processed by the new correction method. We keep all the data types of inputs and outputs of each module the same as HiC-Pro for user-friendliness, and the Long Range Contact Correction module outputs BAM files, in accordance with output file type of the original Read Alignment module.

**Figure 4:**
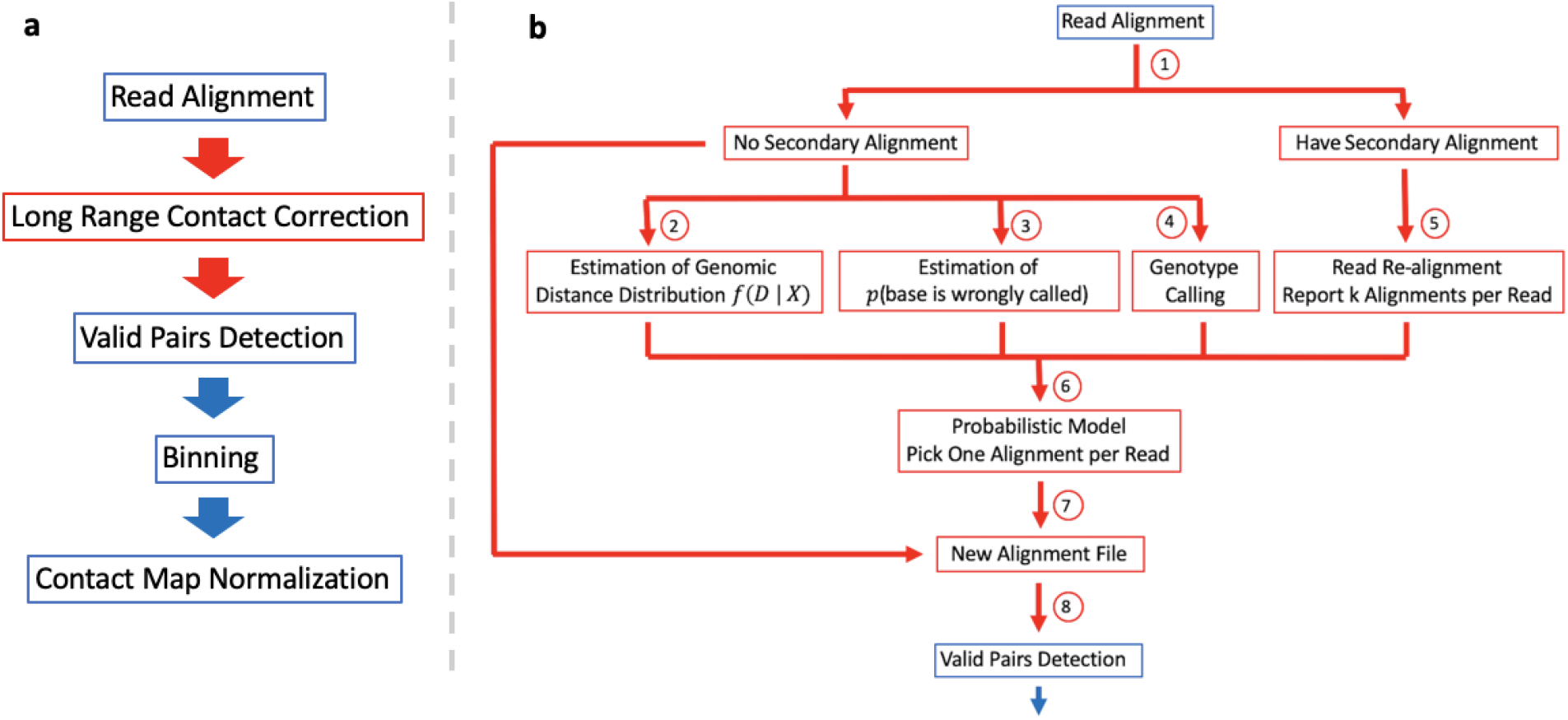
(a) LLC-HiC workflow. Reads are first aligned to the reference genome. Alignment results are then corrected by Long Range Contact Correction module. Invalid pairs such as unmapped reads are discarded in the Valid Pairs Detection module, and the remainings are used for the following Binning and Normalization modules. Blue parts are modules from standard HiC-Pro and the red one is our correction method. (b) The detailed workflow of the module Long Range Contact Correction(red part).

Figure 4b describes the detailed workflow of the module Long Range Contact Correction. After raw read pairs are first aligned, they are separated into two parts (step 1). The first part are read pairs with no secondary alignments, and therefore no need for correction. We call them reliable pairs. The second part are read pairs having secondary alignments in one or both ends that will need correction, we call them ambiguous pairs. We use reliable pairs to estimate the genomic distance distribution *f*(*D* | *X*) (step 2). Similar to Duan et al. [2010], we subdivide genomic distance into discrete bins (0-5kb, 5-10kb, etc.). We assign reliable pairs into these bins according to their alignment results. Then we calculate the bin-level distance distribution *f*(*D* | *X*), which equals the number of pairs in that bin divided by the total number of read pairs.

Reliable pairs are then used for estimating the probability of base calling error ***p***(base is wrongly called) (step 3). Each base is assigned a quality score (QS) after Bowtie2 alignment, which could be used for computing the base calling error by the equation *QS* – 33 = –10log_10_ ***p***(base is wrongly called) [Ewing et al., 1998]. However, it has been shown that these quality scores are inaccurate [Cabanski et al., 2012, DePristo et al., 2011, Li et al., 2009], hence they need to be recalibrated. We design a way to estimate base calling error empirically (step 3). For each quality score *M*, we count the total number of bases with quality score *M* and compute the percentage of these bases that have mismatches with the reference, that is:

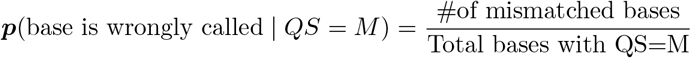

For each base in one read, *r*[*i*], we have

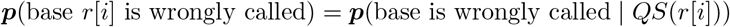

where *QS*(*r*[*i*]) is the quality score of *r*[*i*].

Reliable pairs are also used for genotype calling (step 4), for which we use Genome Analysis Toolkit (GATK) variant calling pipeline [Van der Auwera et al., 2013].

We use Bowtie2 to do read re-alignment of ambiguous pairs. For each read, more than one alignment locations are reported, composing sets of possible alignment choices for the subsequent probability calculation.

Results from step 2 to step 5 are then used for the probabilistic method described in section 3.1 (step 6). In step 6, ambiguous read pairs are assigned to new alignment locations according to posterior probabilities, and these results are finally combined with reliable pairs (step 7) to create new alignment BAM files (step 8) which are used for the following Valid Pair Detection module. Note that step 2 to step 5 can be parallelly computed, therefore our new method does not take much more time than the original HiC-Pro pipeline.

## 4 Results

### 4.1 LLC-HiC can effectively remove erroneous contacts while preserving real signals

We experimented with two variants of the LLC-HiC pipeline, LLC-HiC without genetic variants information (denote it as LLC-HiC in the following context) and with genetic variants information (LLC-HiC-GC), on all four cell types. Both of these two new pipelines are able to remove erroneous contacts effectively. Figure 5a shows the same heatmap region as Figure 1. After using LLC-HiC or LLC-HiC-GC, conserved contact signals in this region are all removed from the heatmap, both in raw data and normalized data (Figure 5a and Figure S8; note that these two figures take cell type IMR90 and KR normalization as an example, but results are the same for all the four cell types).

**Figure 5:**
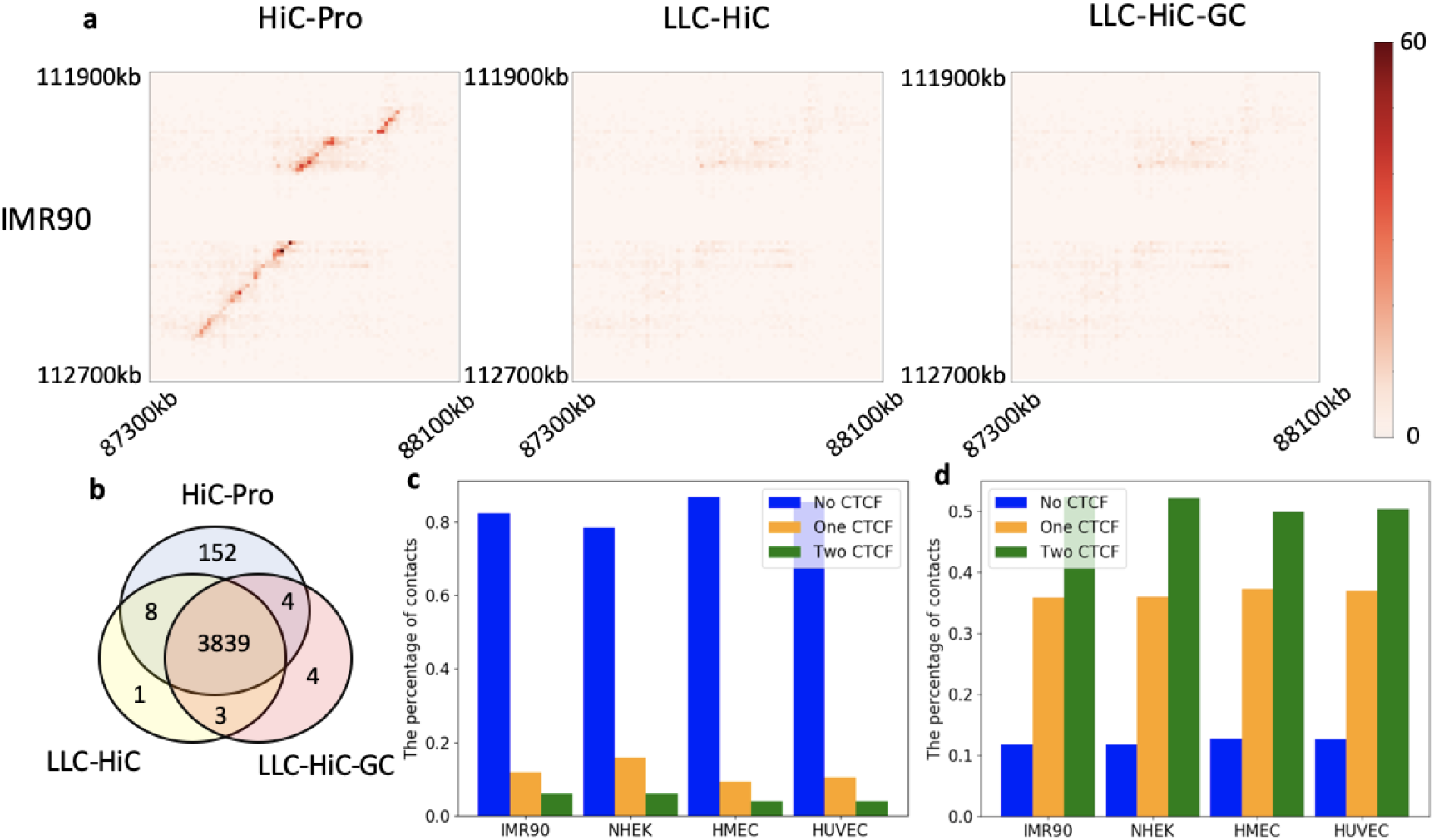
(a) Representative example of comparisons of three methods. Region: chromsome 2 with two genomic intervals 87300 kb-88100 kb and 111900 kb-112700 kb, cell type: IMR90, resolution: 10 kb. Heatmaps are from non-normalized Hi-C matrices. HiC-Pro (the left) is not able to correct these contacts while both LLC-HiC (the middle) and LLC-HiC-GC (the right) remove them successfully. (b), (c) and (d) are Fit-Hi-C results comparison of these three methods. (b) Overlap in conserved contacts of these three methods. (c) The percentage of 152 conserved contacts detected by HiC-Pro uniquely that have zero, one or two bins containing CTCF peaks in different cell types. (d) The percentage of 3839 conserved contacts detected by all these three methods that have zero, one or two bins containing CTCF peaks in different cell types.

We then used Fit-Hi-C again on Hi-C matrices processed by our new method, and compared Fit-Hi-C results on all four cell types to obtain conserved contacts. We then counted the intersection of these conserved contacts from three different pipelines (Figure 5b). LLC-HiC and LLC-HiC-GC have almost the same conserved contacts, and most of these contacts (3839) are also detected from the standard HiC-Pro pipeline. However, there are 152 contacts detected by the standard HiC-Pro which are not detected by either LLC-HiC or LLC-HiC-GC. In every cell type, around 80% of those 152 contacts contain no CTCF peaks in either of two bins (Figure 5c). In comparison, most of those 3839 contacts contain CTCF signals in at least one of two bins (Figure 5d), suggesting that these are more likely to be real contacts.

### 4.2 Comparison of loops detected from three different pipelines

Chromosome loops are usually defined as a cluster of several significant contacts instead of just one entry (one contact signal) in Hi-C matrices [Durand et al., 2016]. Under this definition, many contacts detected by Fit-Hi-C can not be regarded as individual loops. Therefore, we used another widely used loop calling algorithm, HiCCUPS [Durand et al., 2016], to detect loops. We used the same parameters of HiCCUPS as Rao et al. [2014]. The resolution of Hi-C matrices was 10 kb. We compared HiCCUPS results from HiC-Pro, LLC-HiC and LLC-HiC-GC. Similar to the result in Figure 5b, a large number of loops (8528) overlap among all three methods (Figure S9), but there are still 196 loops detected from HiC-Pro uniquely, and overall 92 loops which are newly detected from LLC-HiC or LLC-HiC-GC but not HiC-Pro.

Previous studies showed that besides CTCF, cohesin complex component RAD21 and the structural maintenance of chromosomes protein SMC family are related to the formation of loops [Rao et al., 2014, 2017]. Therefore, we examined the enrichment of CTCF, RAD21 and SMC3 around loop anchors across three categories: loops uniquely from HiC-Pro (196), loops from all three methods (8528), and loops from LLC-HiC or LLC-HiC-GC but not HiC-Pro (92) (Figure 6 and Figure S10). We observed that all three proteins are not enriched around anchors of loops in the first category, but in contrast there are significant enrichment signals appearing around anchors of the 8528 overlapping loops. Furthermore, clear enrichment signals can be observed around anchors of loops in the third category, although not as significant as shared loops. These results give further evidence that loops removed by our method are likely artifacts and our method reassigns read pairs composing those erroneous loops to more likely regions. The reassignment generates new loops that are supported by CTCF, RAD21 and SMC3 enrichment.

**Figure 6:**
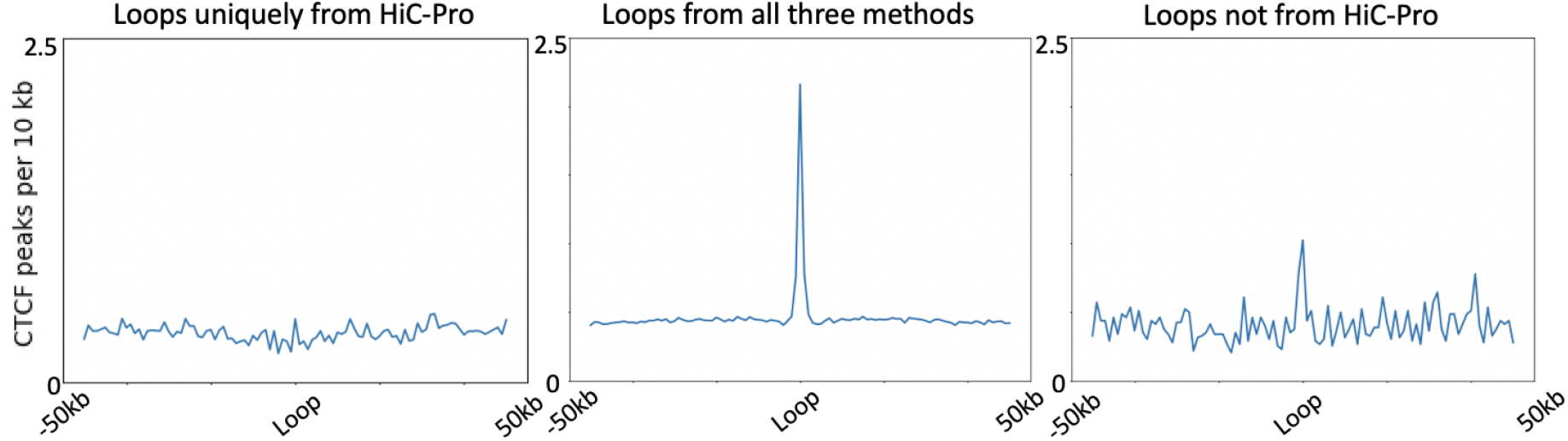
CTCF peaks surrounding loop anchors. cell type: IMR90. The left column: 196 loops that are detected uniquely by the standard HiC-Pro; The middle column: 8528 loops that are detected from all three methods; The right column: 92 loops that are detected from LLC-HiC or LLC-HiC-GC but not HiC-Pro.

### 4.3 LLC-HiC correction on cohesin-independent long-range loops

Rao et al. [2017] detected cohesin-independent long-range loops from auxin-treated HCT-116 cells. By visually examination, they found that some of them were false positives because of structural rearrangements or repetitive regions. We used our LLC-HiC method on auxin-treated HCT-116 Hi-C data to see if we can remove the false positives caused by chromosomal repetitive regions automatically.

We downloaded raw Hi-C reads of auxin-treated HCT-116 cells (GEO: GSE104334), processed them by HiC-Pro or LLC-HiC respectively. Then we called loops by HiCCUPS with the parameters used for annotating super long-range loops as described in Rao et al. [2017] STAR Methods (Resolution: 50 and 100kb; p=2,1; i=4,2; fdr=10%, 10%; d=100 kb, 200 kb). As Rao et al. [2017] did, for both of these two pipelines we additionally filtered out loops that had genomic distance ≤5Mb, had less than a 2-fold ratio between observed counts and expected counts for any of the local expected, or had fewer than 3 contacts that were clustered in one loop. Finally, there were 120 loops detected from the HiC-Pro pipeline and 100 loops from LLC-HiC (Figure 7a). (The number of loops called in our work is approximately but not exact;y the same as the number (88) reported in Rao et al. [2017]. Since the detailed information of those 88 loops was not released, we are not able to compare theirs with ours. The differences may come from different pipelines used for processing the Hi-C data.) The 23 loops called from the HiC-Pro processed data are removed by LLC-HiC. Since cohesin-independent long-range loops are highly related to super-enhancers [Rao et al., 2017], we compared those 23 removed loops with super-enhancer locations of HCT-116 cells, and found that 22 of them do not overlap with HCT-116 super-enhancer locations, while 34% of the other 100 long-range loops overlap with super-enhancer locations. This result indicates that LLC-HiC is able to help detect and remove those erroneous cohesin-independent long-range loops automatically. Figure 7b shows the same heatmap region as Figure 1. Contact signals in this region are clearly weakened after using LLC-HiC.

**Figure 7:**
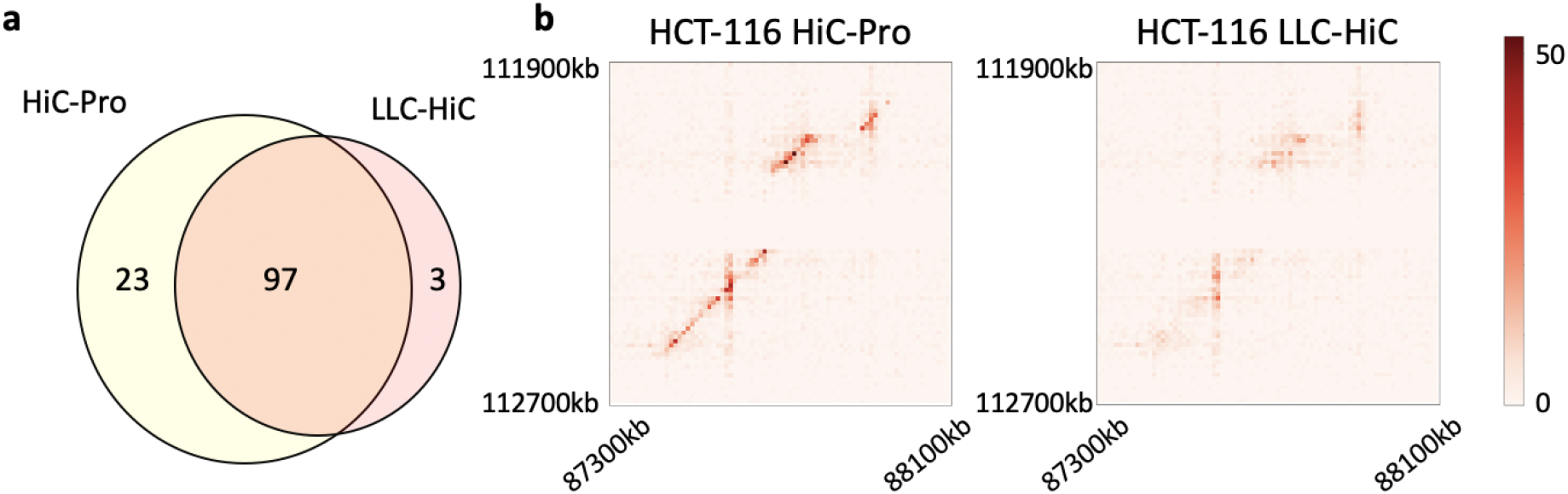
(a) Overlap in long-range loops of auxin-treated HCT-116 cells between HiC-Pro and LLC-HiC. (b) The same region as Figure 1. Celltype: auxin-treated HCT-116, Resolution: 10kb.

### 4.4 LLC-HiC does not significantly alter important features of Hi-C matrices

Our method is designed to filter those erroneous long-range contacts, while keeping other features of Hi-C matrices such as topologically associated domains (TADs) similar to those from previous Hi-C processing pipelines. Thus we used two different measures, HiCRep [Yang et al., 2017] and Jaccard Index (JI), to compare architectures of normalized Hi-C matrices generated from HiC-Pro, LLC-HiC and LLC-HiC-GC.

HiCRep is a method to quantify the similarity between two Hi-C matrices directly, and it returns a score to represent the overall similarity. We ran HiCRep on each pair of intra-chromosomal Hi-C matrices from the same chromosome and the same cell type but different pipelines. For each method pair (for example HiC-Pro and LLC-HiC), we get overall 22 (autosomes) ×4 (cell types) = 88 HiCRep scores. As in Sauerwald et al. [2020], we set the range of a smoothing parameter h required for HiCRep from 0 to 3. According to HiCRep results, Hi-C matrices from three pipelines are highly similar (the mean of all these three distributions are around 0.99), indicating that our method would does not substantially alter on the overall architectures of Hi-C matrices (Figure S11).

We next investigated if our method would alter structures of topological associated domains. Topologically associated domains (TADs) are local chromatin interaction domains in chromatin structures. They are correlated to genome organization and regulation [Dixon et al., 2012]. We used Jaccard Index (JI) to test the similarity between TADs of two chromosomes. JI is a metric measuring the similarity between two sets, which is defined as the size of the intersection of two sets divided by the size of the union of them. We first ran Armatus [Filippova et al., 2014] (gamma-max: 1.0, step size: 0.1) to get TAD sets for each chromosome of all four cell types. Then we used JI on two sets of TAD boundaries as in Forcato et al. [2017] to measure the similarity of TADs. Results reveal that TAD structures are also highly similar among HiC-Pro, LLC-HiC and LLC-HiC-GC, indicating that our method have little impact on TAD structures (Figure S11).

## 5 Discussion

We use a probabilistic method to correct a newly characterised Hi-C artifact by integrating the genomic distance distribution and genetic variations. Our method can remove this new artifact effectively, while preserving original Hi-C features.

One counter-intuitive phenomenon is that integrating genetic variation information in our model does not further improve correction results. Many SNPs leading to incorrect read alignment are not detected by the genotype calling step. Several reasons may cause the failure of genotype calling. First, the coverage of reads from Hi-C data is not high enough for genotype calling, and because of pull down biotin step, most reads will be close to the restriction sites [Khalil et al., 2019], leading to an uneven distribution of reads. In our method, only reads having no secondary alignment are used for genotype calling, which further reduces the coverage. Although adding ambiguous read pairs (pairs having secondary alignment) can increase read depth, incorrect information will be brought in since many such reads are mapped to incorrect locations. We also considered an EM-based algorithm to do correction, however, it is not practical because one round of genotype calling by GATK pipeline is very time consuming. Other information such as read depth signal can be considered in the future, as used in Khalil et al. [2019]. In general, if many reads from one repeat *S*_1_ are aligned to the other repeat *S*_2_, read depth of *S*_1_ will be significantly smaller than the average, and correspondingly, read depth of *S*_2_ will be significantly larger, therefore this character may help improve the accuracy of the remapping step.

Another way to account for genetic variations would be to align Hi-C reads to a genome graph instead of the reference genome. A genome graph is a graph-based data structure which combines the reference genome with genetic variations, hence reflecting genetic diversity of populations [Kim et al., 2019, Ameur, 2019]. We believe that graph-based alignment will increase the accuracy of read alignment and help solve Hi-C artifacts presented in our work. However, a lot of work needs to be done to switch the reference genome to genome graph since many biological algorithms and tools were designed for the reference sequence and are not ready to be compatible with graphs [Ameur, 2019].

Although LLC-HiC is able to correct false positive loops from repetitive regions in cancer cell types such as HCT-116, the method would be invalid for removing erroneous loops caused by structural variations, which is a common phenomena in cancer cells. This would be an interesting direction for the future work.

## 6 Conclusion

We have characterized a previously unstudied artifact from Hi-C data. We show that it is caused by incorrect read alignment due to repeat regions in chromosomes. This kind of artifact leads to long-range chromatin interactions that might be identified as real chromatin loops by mistake. We design a new probabilistic method and combine it with an existing Hi-C processing pipeline, HiC-Pro. We show that our new method is able to filter these erroneous interactions successfully and does not destroy other typical features of Hi-C matrices such as TADs. With parallelization, our method does not take significantly more time than the original HiC-Pro pipeline.

## Acknowledgements

The authors would like to thank Natalie Sauerwald for useful discussion about this work and comments on the manuscript.

---

**Figure S1:**
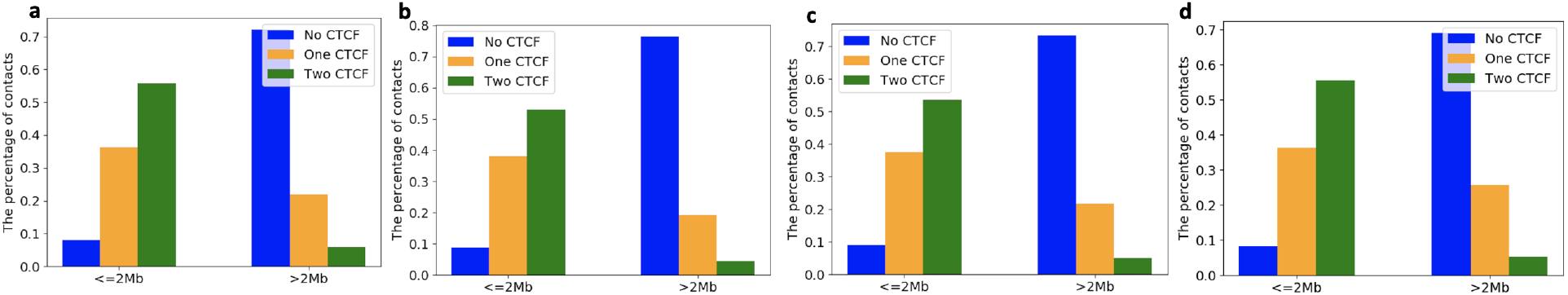
The percentage of conserved contacts with zero, one or two bins containing CTCF peaks of all four cell types, IMR90 (a), HMEC (b), HUVEC (c), and NHEK (d). In each Figure, the left column represents contacts with genomic distance ≤ 2Mb apart, and the right column represents contacts with genomic distance > 2Mb apart.

**Figure S2:**
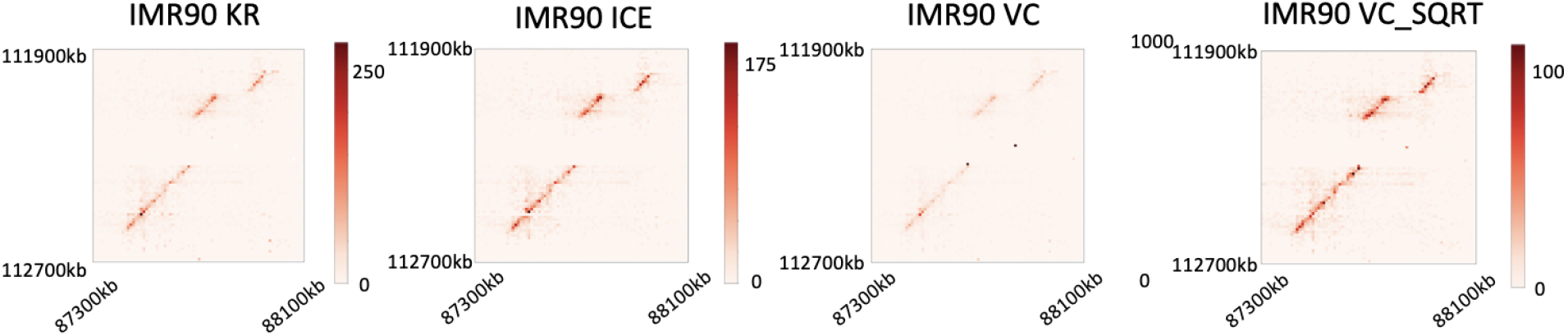
Representative example of a heatmap region containing highly conserved long-range contacts, i.e. two genomic intervals: 87300kb-88100kb and 111900kb-112700kb in chromosome 2. Resolution:10 kb. Heatmaps are from Hi-C matrices of IMR90 with different kinds of normalization.

**Figure S3:**
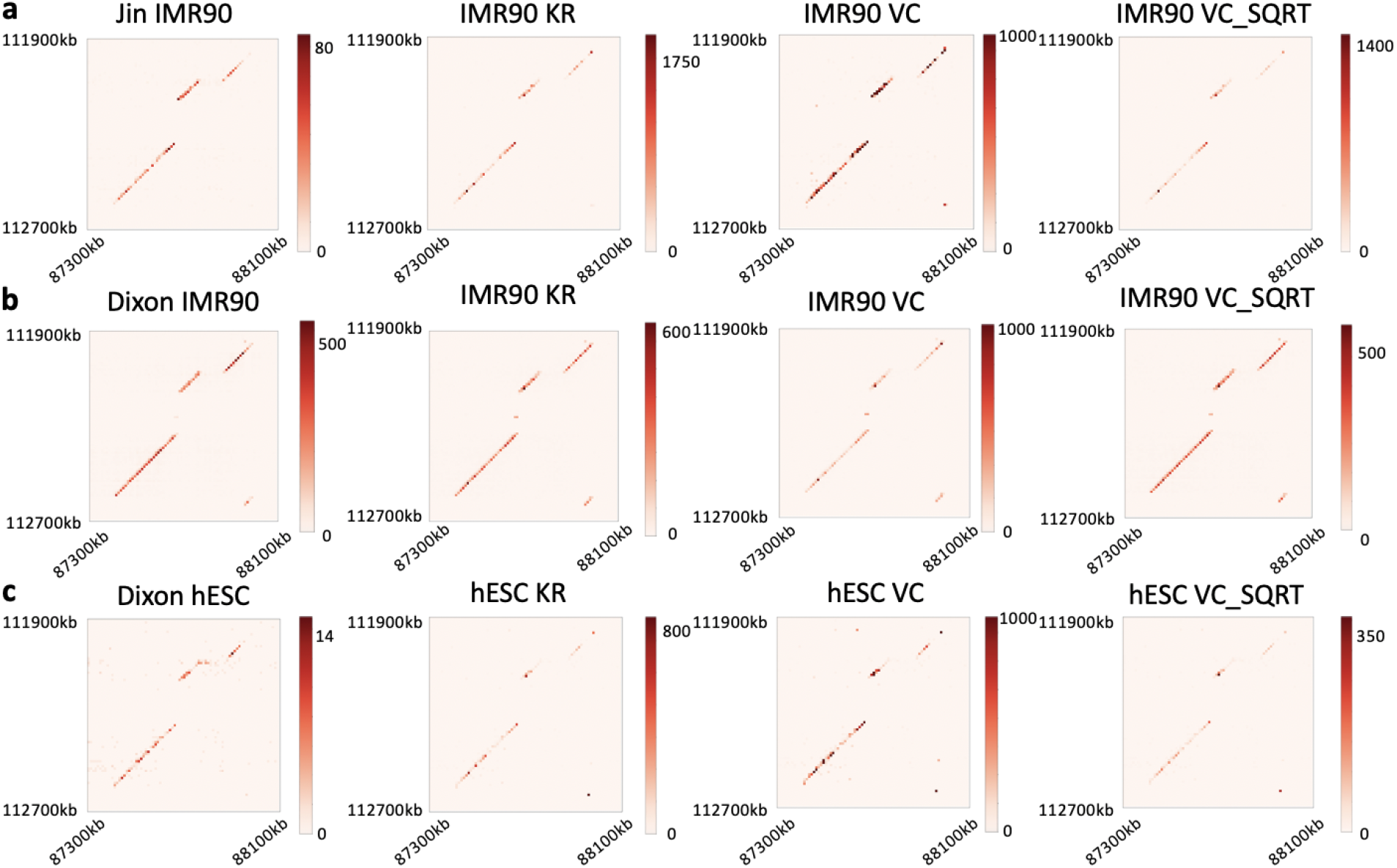
Representative example of a heatmap region containing highly conserved long-range contacts, i.e. two genomic intervals: 87300kb-88100kb and 111900kb-112700kb in chromsome 2. Resolution:10 kb. Data were downloaded from Juicer datasets (http://hicfiles.tc4ga.com/juicebox.properties). Different from Rao et al. [2014], which used in situ Hi-C protocol with restriction enzyme MboI, Hi-C data here were all generated by dilution Hi-C protocol with restriction enzyme HindIII. (a) Heatmaps from the non-normalized as well as three normalized Hi-C matrices of IMR90. Raw reads were from Jin et al. [2013], and were processed by Juicer. (b) Heatmaps from the non-normalized as well as three normalized Hi-C matrices of IMR90. Raw reads were from Dixon et al. [2012], and were processed by Juicer. (c) Heatmaps from the non-normalized as well as three normalized Hi-C matrices of hESC. Raw reads were from Dixon et al. [2012], and were processed by Juicer.

**Figure S4:**
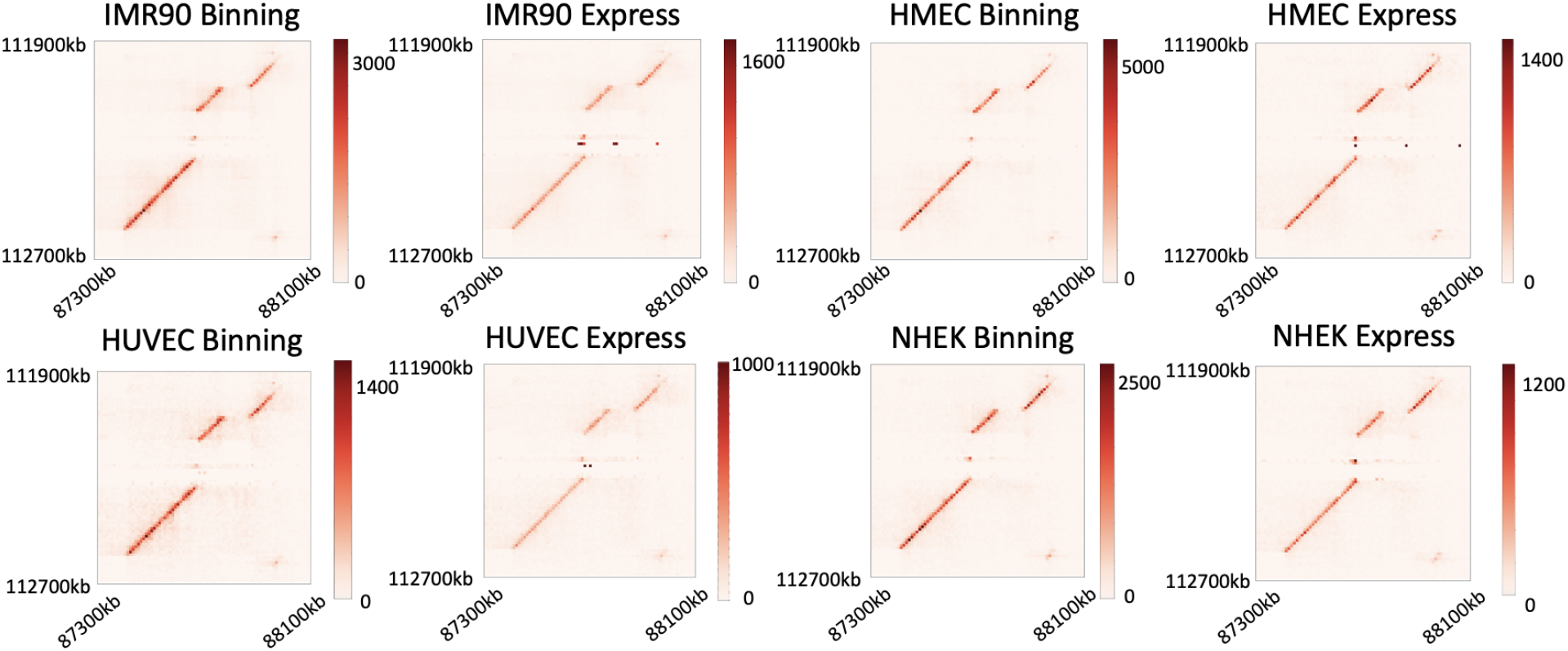
Representative example of a heatmap region containing highly conserved long-range contacts, i.e. two genomic intervals: 87300kb-88100kb and 111900kb-112700kb in chromosome 2. Resolution:10 kb. Raw Hi-C reads of four cell types were from Rao et al. [2014], aligned by Bowtie2, and then processed by HiFive [Sauria et al., 2015]. Here “Binning” and “Express” are two normalization algorithms in HiFive.

**Figure S5:**
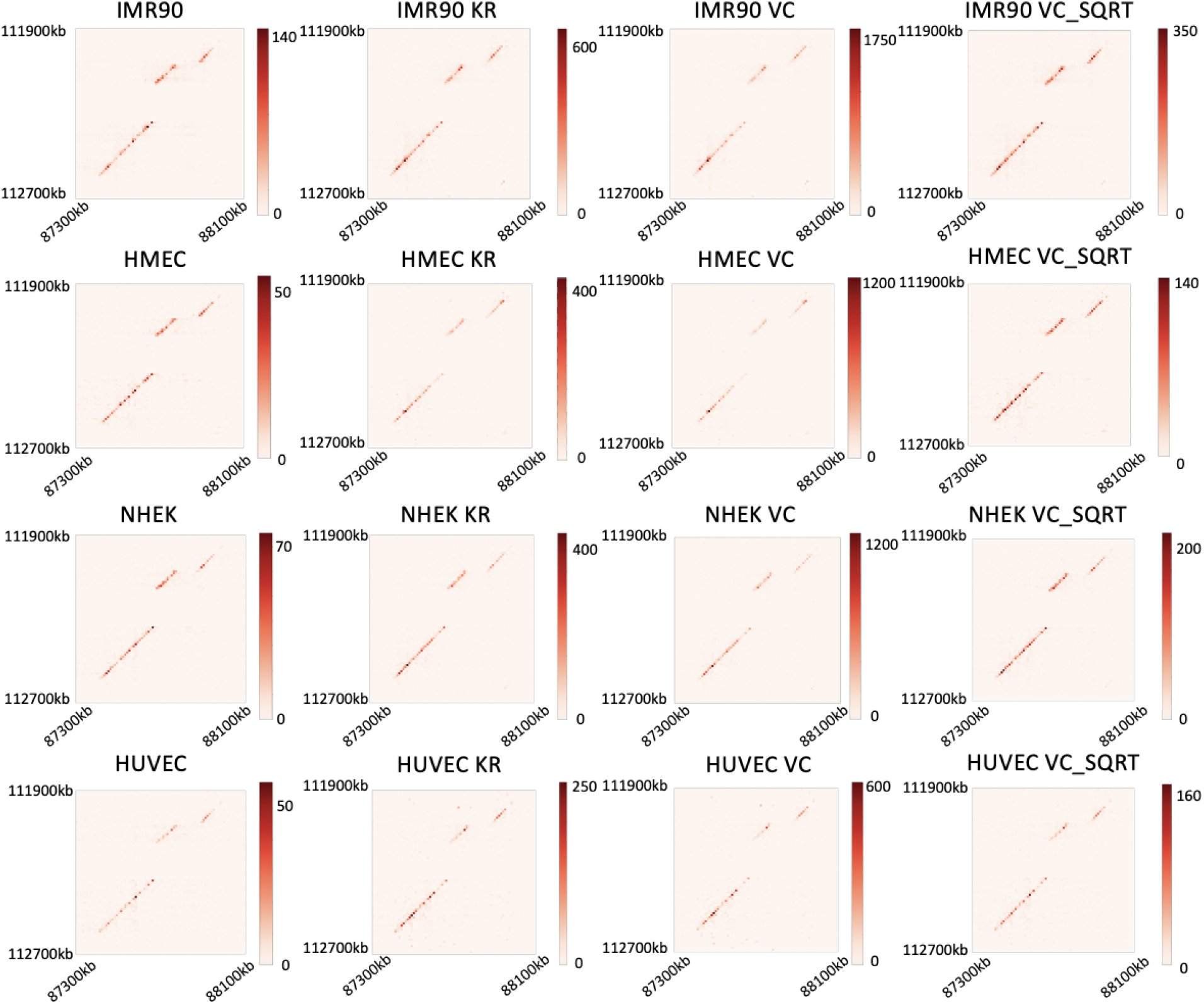
Representative example of a heatmap region containing highly conserved long-range contacts, i.e. two genomic intervals: 87300kb-88100kb and 111900kb-112700kb in chromsome 2. Resolution:10 kb. Raw Hi-C reads of four cell types were from Rao et al. [2014], aligned by BWA [Li and Durbin, 2010], and then processed by Juicer. The first column are heatmaps from the non-normalized Hi-C matrices, and the remainings are heatmaps from normalized Hi-C matrices with different kinds of normalization. Data were downloaded from Juicer datasets (http://hicfiles.tc4ga.com/juicebox.properties).

**Figure S6:**
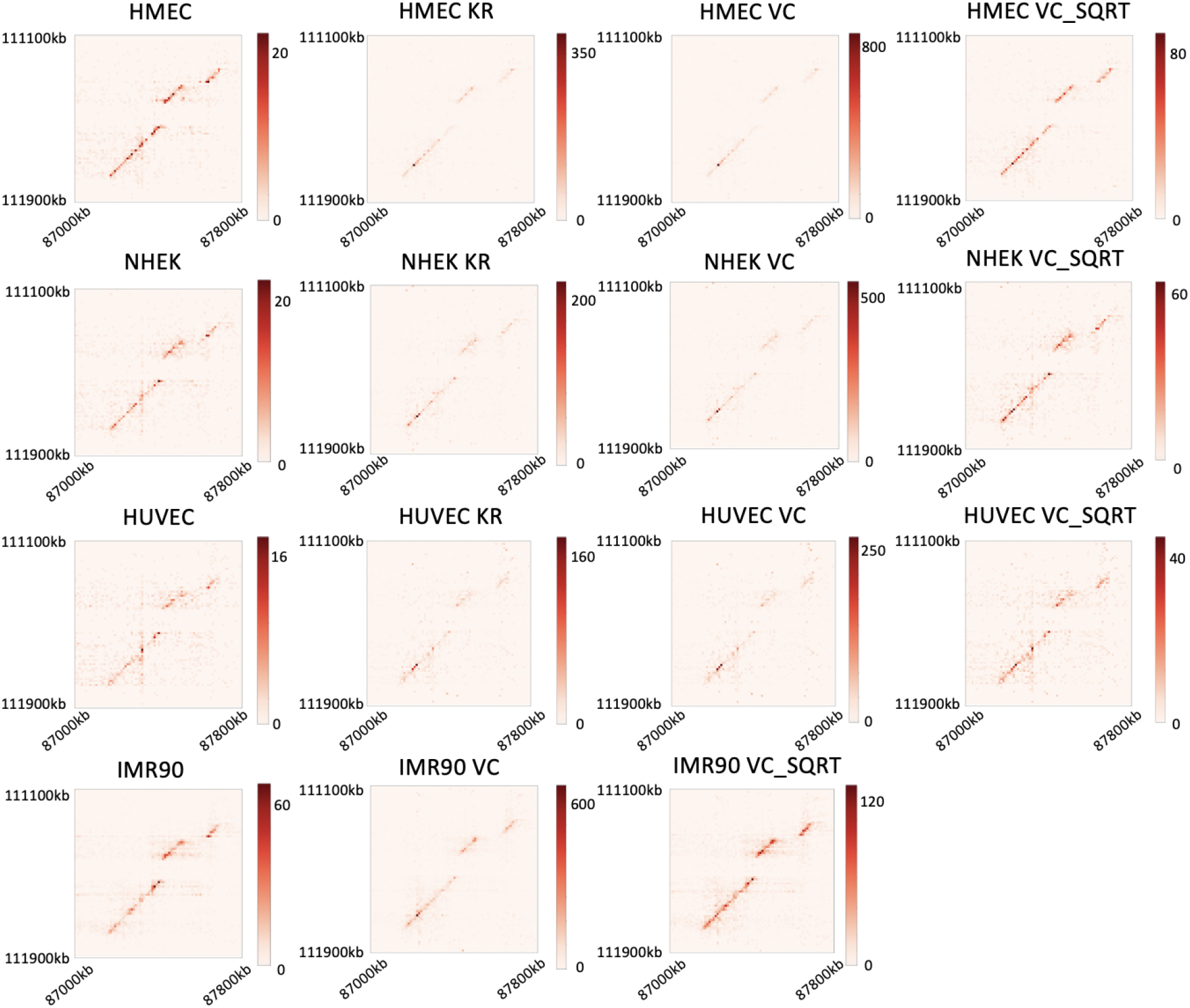
Representative example of a heatmap region containing highly conserved long-range contacts. Raw Hi-C reads of four cell types were from Rao et al. [2014], aligned by Bowtie2, however, unlike Figure 1 which used hg19 as the reference genome, here we used hg38. Note that there is a shift between coordinate system of hg19 and hg38, therefore, two genomic intervals here are 87000kb-87800kb and 111100kb-111900kb in chromsome 2, which is almost the same region as shown in Figure 1. Aligned reads were further processed by HiC-Pro (without ICE normalization), and normalized by Juicer. Here we omit the heatmap of IMR90 cell with KR normalization method since it failed in this case (most values of bins are NaN.)

**Figure S7:**
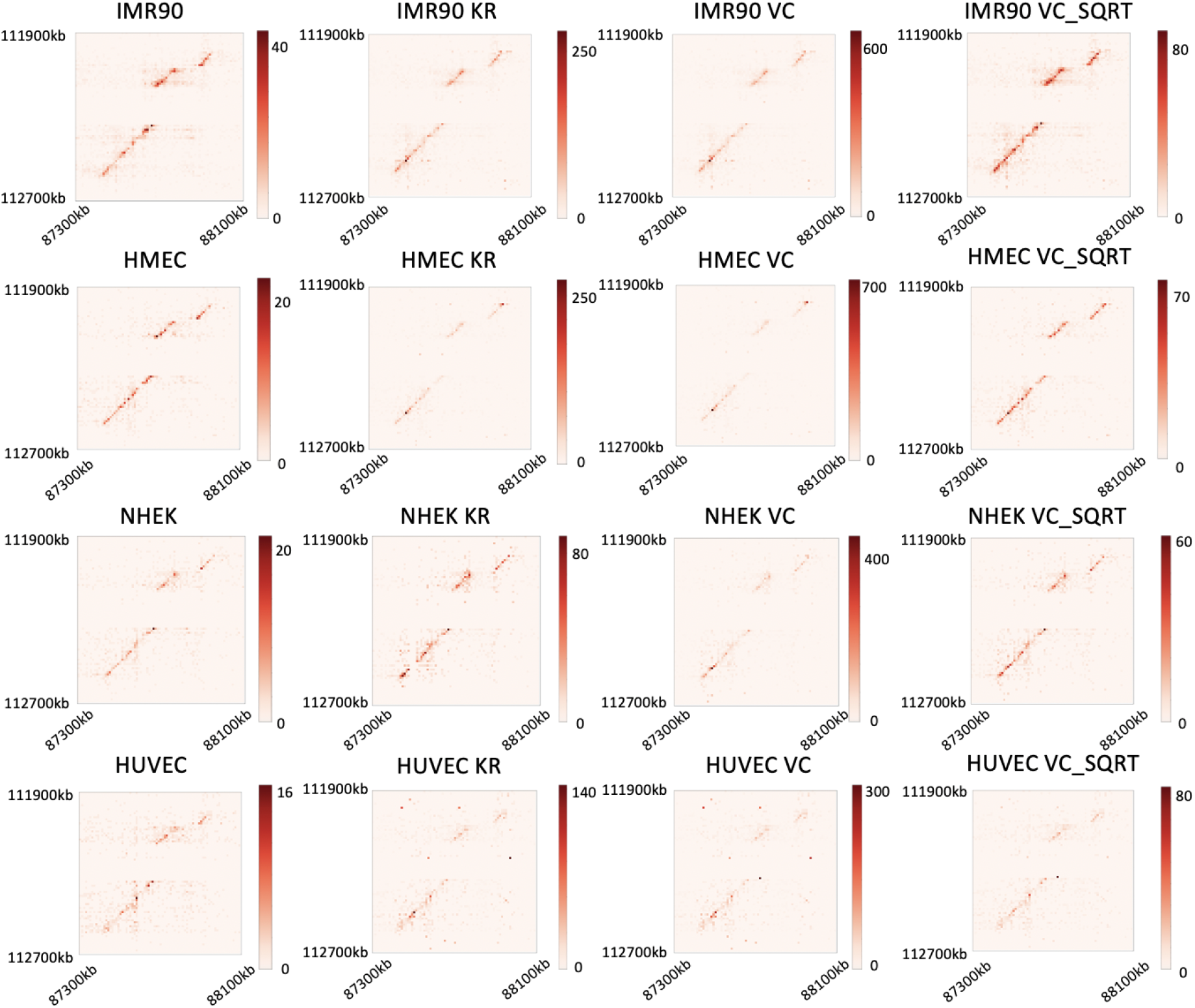
The same region as Figure 1. Raw Hi-C reads of four cell types were from Rao et al. [2014], and processed by HiC-Pro, except that all reads with MAPQ=0 are filtered out.

**Figure S8:**
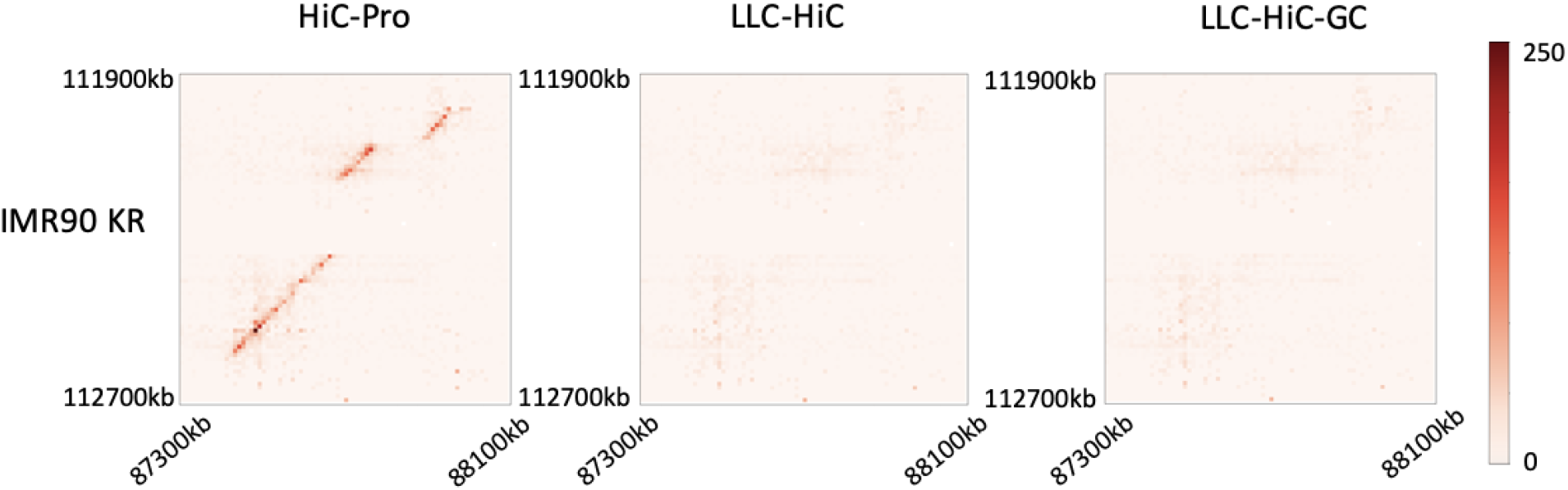
(a) Representative example of comparisons of three methods. Region: chromsome 2 with two genomic intervals 87300 kb-88100 kb and 111900 kb-112700 kb, cell type: IMR90, resolution: 10 kb. Heatmaps are from KR-normalized Hi-C matrices. HiC-Pro (the left) is not able to correct these contacts while both LLC-HiC (the middle) and LLC-HiC-GC (the right) remove them successfully.

**Figure S9:**
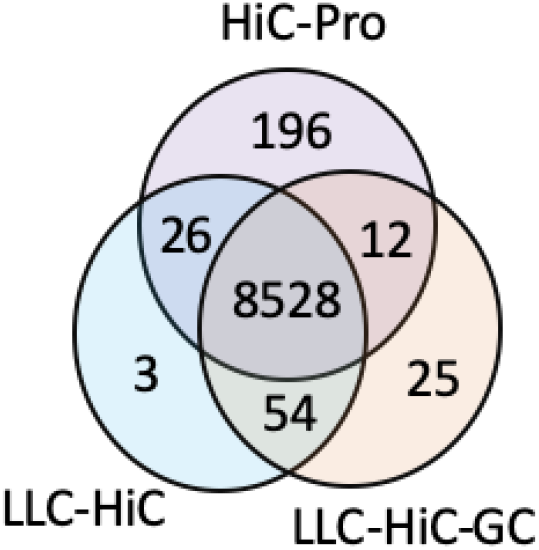
Overlap in detected loops among HiC-Pro, LLC-HiC, and LLC-HiC-GC. cell type: IMR90.

**Figure S10:**
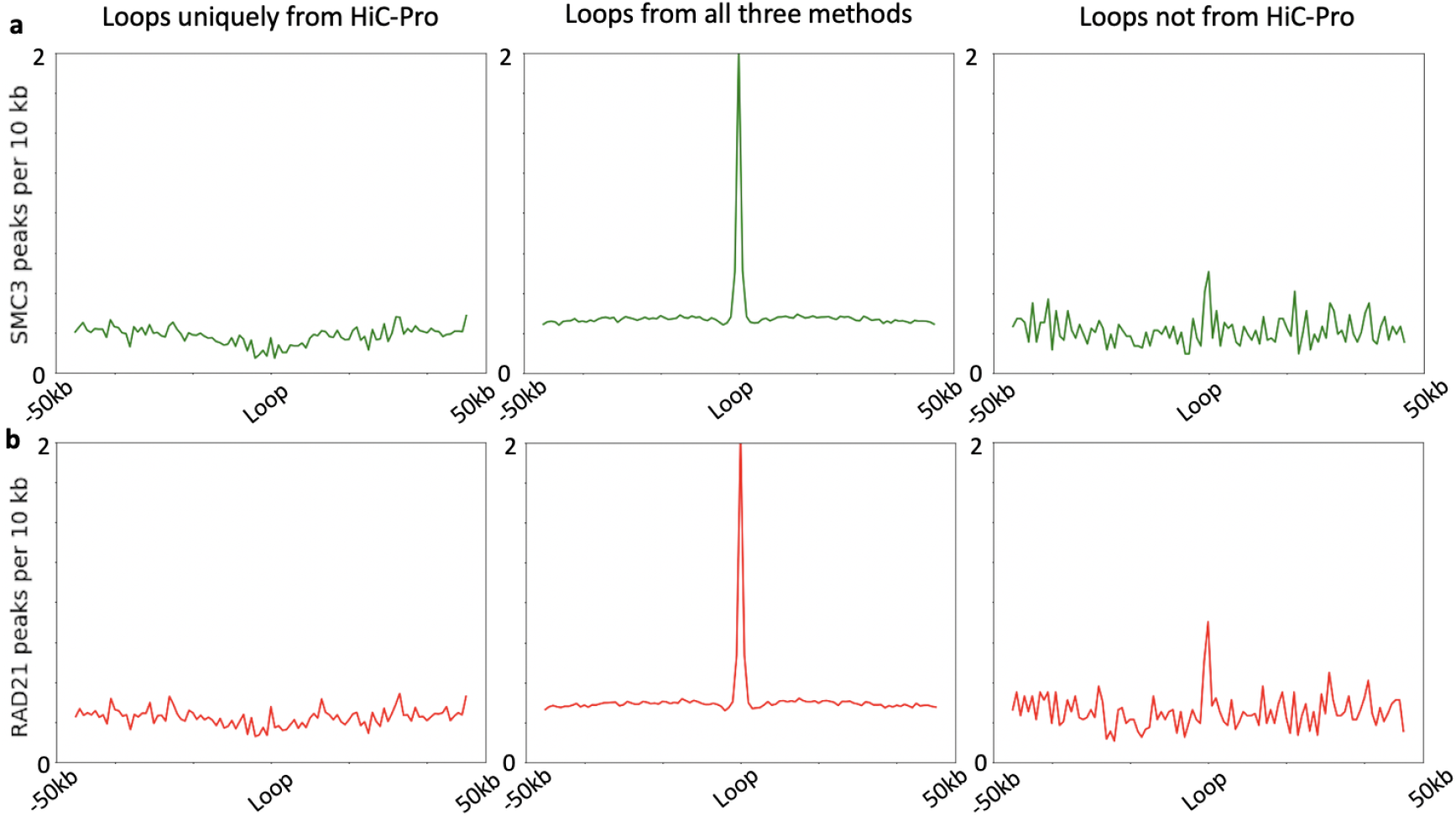
SMC3 and RAD21 peaks surrounding loop anchors. cell type: IMR90. The left column: 196 loops that are detected uniquely by the standard HiC-Pro; the middle column: 8528 loops that are detected from all three methods; the right column: 92 loops that are detected from LLC-HiC or LLC-HiC-GC but not HiC-Pro. (a) SMC3 peaks surrounding loop anchors in three categories. (b) RAD21 peaks surrounding loop anchors.

**Figure S11:**
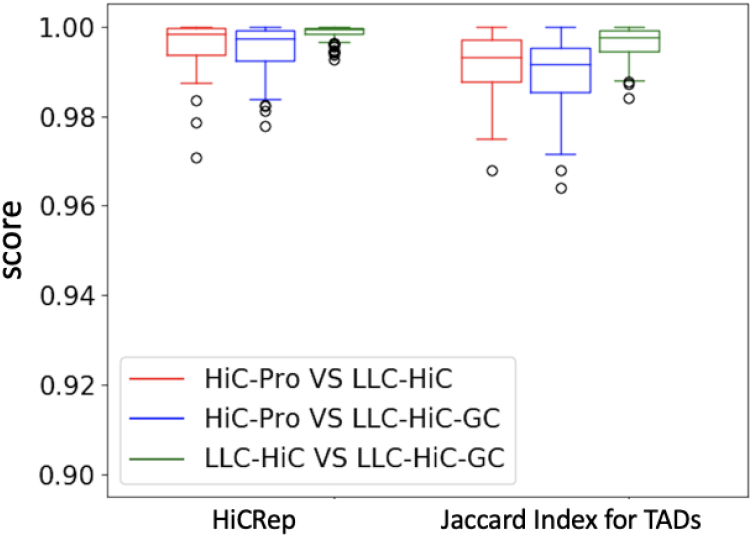
Box plots of HiCRep scores (the left column) and Jaccard Index scores (the right column) between different Hi-C processing pipelines. The higher score indicates higher similarity, and the score value 1 means exact the same.

